# Epidermal cell surface structure and chitin-protein co-assembly determine fiber architecture in the Locust cuticle

**DOI:** 10.1101/793992

**Authors:** Sanja Sviben, Oliver Spaeker, Mathieu Bennet, Marie Albéric, Jan-Henning Dirks, Bernard Moussian, Peter Fratzl, Luca Bertinetti, Yael Politi

**Author notes:** These authors contributed equally to the paper (Oliver Spaeker, Mathieu Bennet). B CUBE – Center for Molecular Bioengineering, Technische Universität Dresden, Dresden, Germany.

## Abstract

The geometrical similarity of helicoidal fiber arrangement in many biological fibrous extracellular matrices, such as bone, plant cell wall or arthropod cuticle, to that of cholesteric liquid mesophases has led to the hypothesis that they may form passively through a mesophase precursor rather than by direct cellular control. In search of direct evidence to support or refute this hypothesis, here, we studied the process of cuticle formation in the tibia of the migratory locust, *Locusta migratoria*, where daily growth layers arise by the deposition of fiber arrangements alternating between unidirectional and helicoidal structures. Using FIB/SEM volume imaging and scanning X-ray scattering, we show that the epidermal cells determine an initial fiber orientation from which the final architecture emerges by the self-organized co-assembly of chitin and proteins. Fiber orientation in the locust cuticle is therefore determined by both active and passive processes.

## Main text

The orientation of fibers in extracellular matrices (ECM), skeletal and connective tissues strongly affects their physical properties, which in turn determines morphogenesis, cell adhesion and functionality(1). Fiber orientation therefore needs to be spatially and temporally controlled by the tissue that produces the ECM (e.g. refs 5-7). For many biological systems it is however still unclear how the control over fiber orientation is achieved. Biological fibrous materials, such as bone, plant cell wall, and arthropod cuticle, are often described as liquid crystal (LC) analogues as they exhibit geometrical similarity to LC, but are solid in their functional state(2). In unidirectional fiber arrangement (analogous to nematic LC) sheets made of parallel fibers are stacked on top of each other with the same orientation, whereas in the helicoidal arrangement, also termed twisted plywood or Bouligand structure (analogous to cholesteric LC) the orientation of the fibers in successive sheets is continuously rotating(2). Both types of fiber orientations can be found in arthropod cuticle made of α-chitin fibers embedded in a protein matrix. They are also present in plant cell walls made of cellulose fibers embedded in a mixed matrix of hemicellulose and lignin, and in mineralized or unmineralized collagen based tissues in vertebrates(2–4).

Furthermore, many biomacromolecules and biological crystals, amongst them, hemicellulose, cellulose, chitin, collagen and silk have been demonstrated to form LC phases *in vitro* (reviewed in (5)). These observations have led to the hypothesis that fibrous biological materials organize by molecular self-assembly via a liquid mesophase precursor that subsequently solidifies by dehydration and cross-linking(2, 6–8). Self-assembly is thought to be facilitated by matrix components such as hemicellulose in the case of plant cell wall or proteins in the arthropod cuticle(3). The latter is supported by molecular genetics studies in several insects species showing modified fiber organization, often with lethal phenotypes, following a genetic knockdown or silencing of certain cuticular proteins(9–13).

In contrast, evidence for cell-directed fiber organization and for the transfer of mechanical stresses from the cytoskeleton into the ECM, as observed in fibroblasts tissue culture(14), calls the general self-assembly hypothesis into question. In an attempt to reconcile self-assembly and cellular control mechanisms for ECM deposition, Bouligand(15) suggested that cell membranes could create strong boundary conditions that influence fiber organization and self-assembly. Neville(16) suggested a two-model system with varying degree of cellular control in the deposition of unidirectional and helicoidal architectures in insect cuticle based on the bilateral symmetry of the former and bilateral asymmetry (the helicoids are always left handed) of the latter. How this is achieved, is the subject of this work. We studied the circadian clock regulated cuticle deposition in the locust, *Locusta migratoria*. In this species, endocuticle formation involves alternating deposition of non-lamellate layers of unidirectional fiber arrangement during the day and lamellate cuticle made of twisted plywood structure during the night. (Fig. 1)(17). By means of 2D and 3D machine learning analysis of focused ion beam – scanning electron microscopy (FIB/SEM) volumes and high-resolution scanning X-ray scattering methodologies, we show that epidermal cell surface structure and chitin-protein co-assembly determine fiber architecture in the Locust cuticle.

**Fig. 1.**
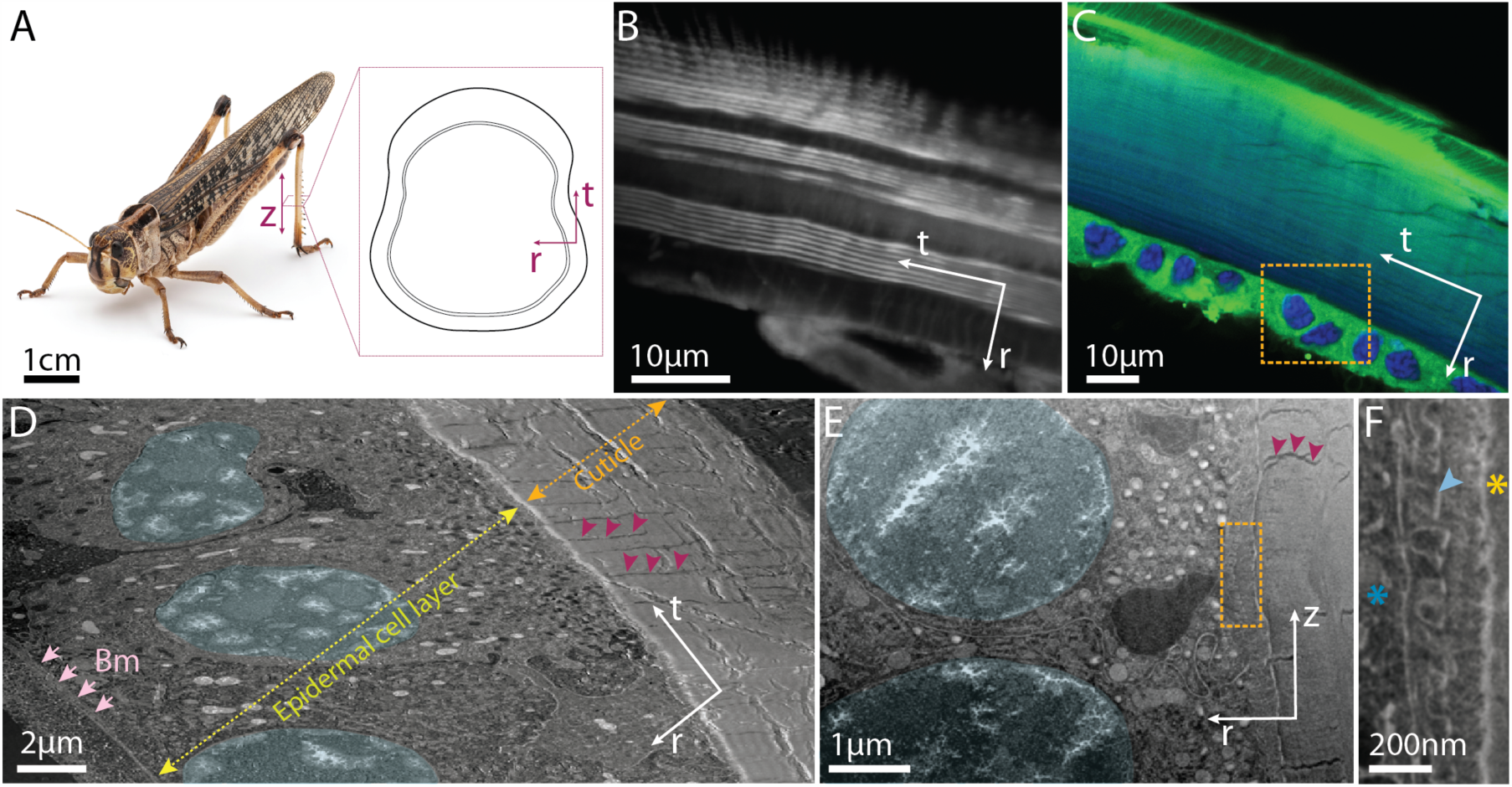
Cuticle deposition in *Locusta migratoria*. (A) Adult locust. Inset showing a cross section of the hind-tibia and coordinate system used throughout: (*z*) longitudinal, (*r*) radial and (*t*) transverse directions. (B) Confocal Light Scanning Microscopy (CLSM) image of tibia cross-section stained with Direct Yellow 96 stain showing the daily growth layers of chitin fibers in unidirectional and helicoidal fiber arrangements, leading to respectively alternating non-lamellate and lamellate layers. (C) CLSM tibia cross-section double-stained with DAPI (blue) and Nile red (green) showing the epidermal cell layer adjacent to the cuticle. Orange rectangle indicates similar area of the cross-section imaged using FIB/SEM in (D). (D) SEM micrograph obtained by serial slice-and-view using FIB/SEM showing epidermal cells between the basal membrane (Bm) and the cuticle. The nuclei are false colored in light blue and the pore canals are indicated with magenta arrowheads. (E) SEM micrograph obtained by re-slicing 3D FIB/SEM volume showing the assembly zone between the cuticle (right) and the surface of the epidermal cell layer (left). Magenta arrowheads point to a pore canal, cell nuclei are false colored in light blue. The orange rectangle marks an area from which the micrograph shown in (F) is taken. (F) Magnification of the assembly zone in a region indicated in (E). The micrograph is obtained from a different depth in *t* direction than shown in (E). The microvillar structures on the apical cell surface which end with bright contrast at the location of the presumably chitin synthesizing plaques (light blue arrowhead), can be seen as well as the newly deposited cuticle (yellow asterisk). Cell interior is marked with blue asterisk.

## Results and discussion

The innermost part of the procuticle of *L. migratoria*, termed endocuticle, is deposited by a single layer of epidermal cells that secrete the cuticle’s components into a 0.5-1 µm wide “assembly zone” (Fig. 1C-F)(18). Chitin is synthesized and secreted by a transmembrane enzyme, chitin synthase, located within “plaques” at the tips of microvilli emerging at the apical cell surface (Fig. 1F). Cuticular proteins are proposed to be delivered to the assembly zone by vesicles(18). We imaged the deposition zone in hind tibia cuticle collected during either “day” or “night” conditions (hereafter termed *Day* or *Night* samples, respectively) using FIB/SEM volume imaging in order to determine the spatial relationships of the microvilli with respect to each other, to the fibers within the assembly zone and within the cuticle, and to the cuticle surface (Fig 2). For segmentation and quantification we implemented a three-dimensional machine learning (3D-ML) algorithm using a U-shaped, 3D fully convolutional networks as proposed recently(19). Cryofixed resin embedded *Day* and *Night* samples show differences in the microvilli structure and organization (Fig. 2). In *Night*, we mainly observed individual microvilli structures containing a single plaque, but occasionally no more than two microvilli are seen merged (Fig. 2A-D). In *Day*, the bases of 2-4 microvilli, each containing a single plaque at its tip, are merged to form ridges of around 300 nm (Fig. 2H-K, Supplementary Information Fig. 1).

**Fig. 2.**
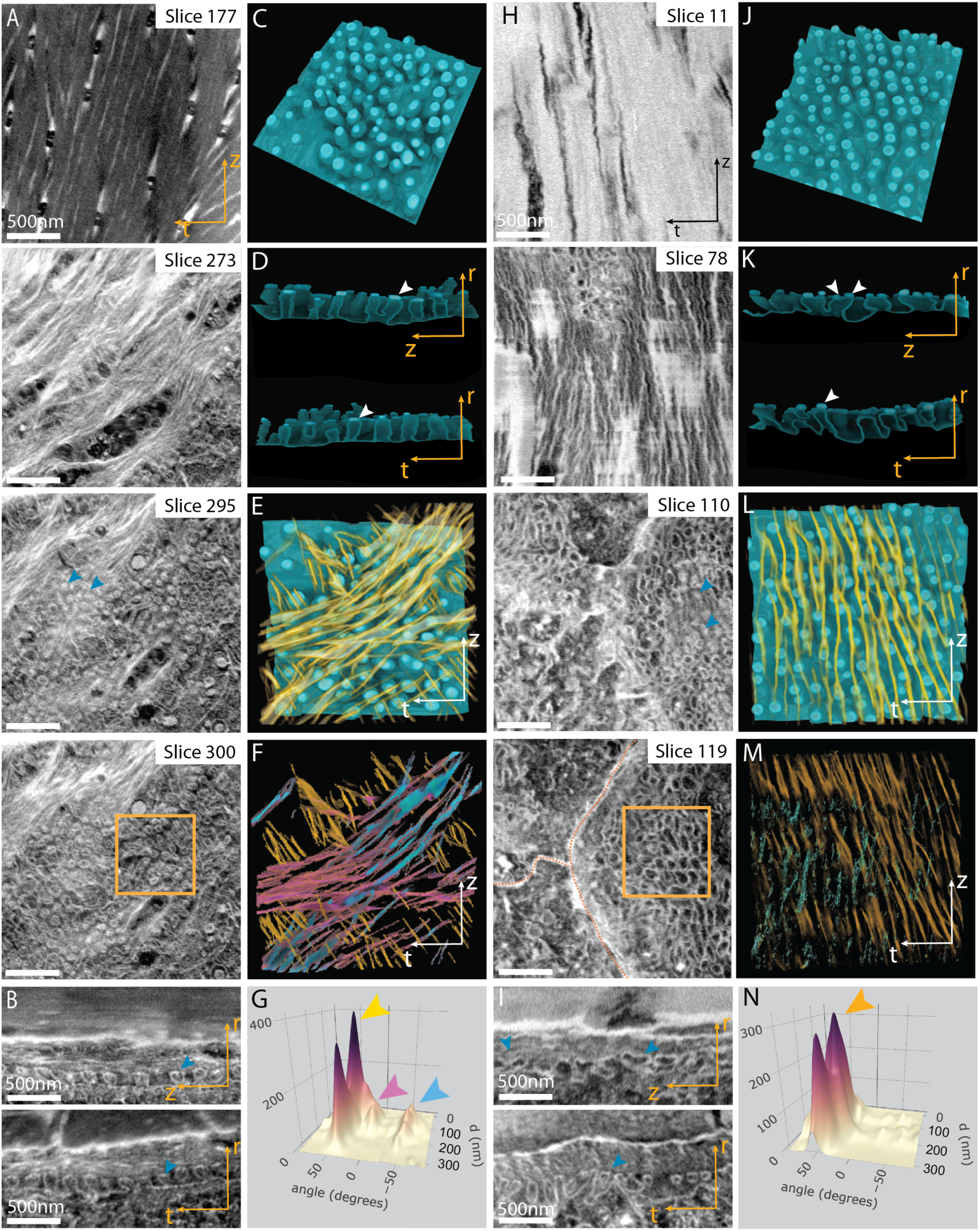
Fiber deposition within the assembly zone. Reconstruction and quantification of 3D FIB/SEM data obtained from cryo-fixed locust tibiae samples. (A, H) Series of FIB/SEM slices (along the *r* direction from the cuticle to the epidermal cell surface) from *Night* (a) and *Day* (H) samples. Cyan arrowheads point to plaques at the tips of the microvilli. Orange rectangles in (a) and (H) indicate the regions for which 3D volume rendering are shown in (C-F) and (J-M), respectively. (B, I) FIB/SEM images obtained by re-slicing the FIB/SEM 3D datasets along *(t)* (top) and *(z)* (bottom) directions, for *Night* (B) and *Day* (I) samples. The comparison shows that the microvilli have similar dimensions in both *(z)* and *(t)* directions in the *Night* samples (b, top and bottom) whereas they are elongated along the (*z*) direction in the *Day* samples (I, top). (C, J) Volume rendering of the apical surface of the epidermal cells in *Night* (A) and *Day* (J) samples. The plaques are depicted in bright blue. (D, K) *(zr)*, and *(tr)* plane views of resliced volumes. Only one plaque (white arrowhead) is situated at the tips of each microvillus in *Night* samples (D), but 2 or 3 plaques are observed in the *Day* samples on top of microvilli merged along the *(z)* direction (K). (E, L) 3D volume rendering of the chitin fibers/fiber bundles (yellow) observed in the assembly zone and the apical cell surface in *Night* (E) and *Day* (L) samples. For simplicity only half of the assembly-zone thickness is shown. (F, M) Orientation color-maps of the fibers in the assembly zone (entire volume) in *Night* (F) and *Day* (M) samples. (G, N) Fiber orientation (angle) vs. assembly zone depth (D) histograms showing the variation in fiber orientation as a function of their position within the assembly zone along the *(r)* direction from the cell-surface (0 nm) to the cuticle (300 nm). The color (yellow, pink, blue, orange) arrowheads represent the respective dominant orientations in (F) and (M).

In order to determine the microvilli organization over large areas (hundreds of microns) we prepared samples using traditional chemical fixation methods and osmium staining (see methods for details) followed by resin embedding. These samples are overstained and the images contain less details (Supplementary Information Fig. 2), which facilitates automatic image analysis using two-dimensional machine learning (2D-ML) networks(20). In both *Day* and *Night* samples cell surface regions containing microvilli extend over several cells, but the spatial organization of the microvilli differs substantially between the two conditions (Supplementary Information Fig. 2, 3 C and D) which is in agreement with the results obtained with cryofixed samples. A comparison between the 3D Fourier transform (FT) of labeled volume of the microvilli in *Day* and *Night* datasets (Supplementary Information Fig. 3 E-H), shows the differences in microvilli packing and organization. In *Night* samples, the organization is short ranged whereas in *Day* samples we observe second order correlations indicating long-range order. Furthermore, the arrangement is highly anisotropic in *Day* samples, where the distance (*d*_*L*_) between the center of the ridges determined from the FT, is 140 nm laterally, and 250 nm vertically (*d*_*V*_) but only slightly anisotropic in the *Night* samples (*d*_*L*_ = 140 nm, *d*_*V*_ = 180 nm). During the de-novo formation of the embryo cuticle in *Drosophila melanogaster*, the apical membrane of the epidermal cells forms morphologically similar ridges, the apical undulae. However, in that case they are elongated over several microns(21).

The preservation of the assembly zone in cryofixed resin embedded sections allowed determining the thickness and the orientation of the freshly deposited fibers in 3D. Using FIB/SEM data we determined fiber-bundle thickness of around 20 nm. However, inspecting 100 nm thin sections using Transmitted Electron Detector (TED) in the FIB/SEM we also observed thinner fibers around 5 nm (Supplementary Information Fig. 4). In both *Day* and *Night* samples the fibers/fiber bundles assume parallel orientation with respect to the cuticle and the cell surface. The fibers/fiber bundles are stacked on top of each other (in the radial direction) with a slight lateral shift, giving the false appearance of oblique fibers protruding from the microvilli towards the cuticle when viewed in 2D in the *(rt)* plane (Supplementary Information Fig. 4). We speculate that the microvilli are dynamic structures that move laterally in the *(tz)* plane while depositing the chitin fibers.

The fibers long axes in the *Day* samples are parallel to the microvilli ridges long axes (Fig. 2H and L). In *Night* samples, we observe a narrow distribution of fiber orientation around three dominant orientations that seem to reflect the packing pattern of the microvilli (Fig. 2A and E). In these samples, the dominant orientation of the fibers changes with the “depth” of the assembly zone, i.e. from the cell surface to the cuticle (Fig. 2M), (with ∼50 degree shift). We deduce from this observation that during the dark phase, the fibers are secreted to the assembly zone in discrete orientations (Fig. 2 E-G). However, within the cuticle, the rotation angle between successive sheets is much smaller (in the order of 1 degree). This implies that at night the organization of the fibers to helicoidal structure occurs *via* a self-assembly mechanism that includes re-orientation and compactization of the fibers, where compactization refers to the process of fiber packing in the radial direction from the cell surface to the cuticle-surface. In *Day* samples, as the fibers are deposited in their final orientation, compactization occurs without reorientation.

In addition to the microvilli, in both *Day* and *Night* samples, we often observed large regions displaying different cell surface structures and a large number of vesicles (Fig. 3, Supplementary Information Fig. 5-7). These regions are at least as abundant as regions with microvilli. Occasionally, structures resembling the plaques in shape are present in these regions (Supplementary Information Fig. 7). This cell surface structure shows similarities to that found in kidney epithelial cells during remodeling of microvilli (22). The assembly zone at this stage is narrower and highly stained such that the fibers cannot be discriminated (Supplementary Information Fig. 4B). We speculate that this stage may be related to vesicular protein secretion and potentially to remodeling of the microvilli. Thus, our observations suggest an active cell surface with temporal process with at least two time scales, one in which dynamic microvilli move laterally while depositing chitin fibers and a second stage in which the microvilli disappear and are replaced by vesicles.

**Fig. 3.**
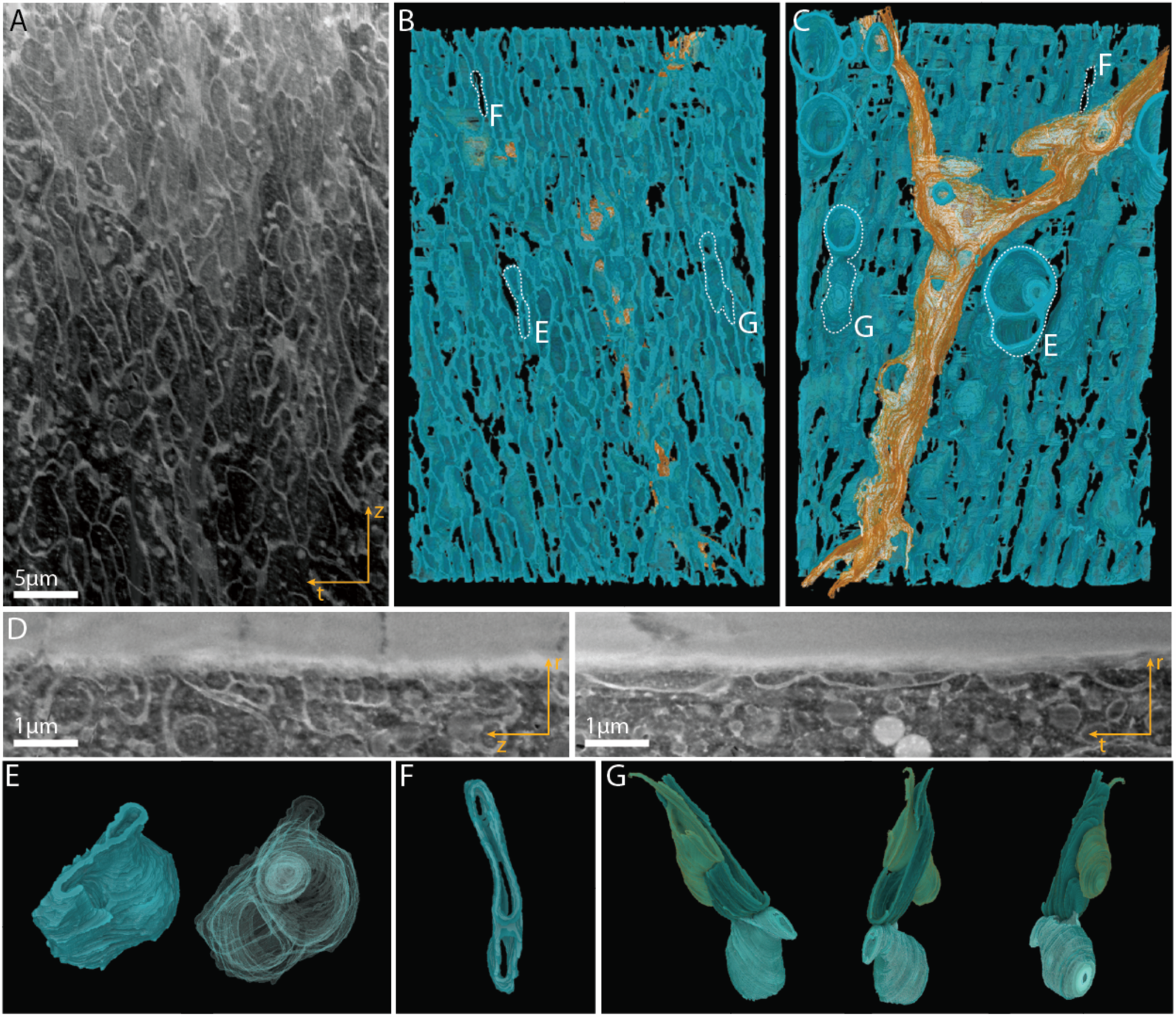
The apical surface of the epidermal cells in the absence of microvilli. (A) FIB/SEM slice at the cell surface showing irregular structures in a *Day* sample. (B, C) Volume rendering of the 3D reconstructed FIB/SEM data represented in (A) in the *(tz)* plane. In (B) the cell surface is viewed from the outside of the cell. In (C) the cell surface is viewed from within the cells outwards. Lateral cell membranes are depicted in orange. Individual structures depicted in (E-G) are marked with a white dash line in both (B) and (C). (D) 3D FIB/SEM data resliced to show the *(tr)* and *(zr)* plane views. (E-G) 3D volume rendering of representative structures visible in (B, C). (E) Multivesicular body. (F) Elongated openings at the cell surface. (G) Vesicles presumably fusing with the apical cell membrane.

3D FIB/SEM is an excellent tool to characterize meso-scale structures like the apical cell surface and the fiber orientation, however this technique is insensitive to the molecular structure and organization of the chitin fiber and associated proteins. In order to gain further insight into the chitin and protein molecular assemblies we used scanning X-ray scattering. Thin sections (∼30 µm) of locust hind tibiae were scanned using a monochromatic focused X-ray beam (beam diameter ∼1 µm) while 2D X-ray diffraction patterns and X-ray fluorescence (XRF) signals were recorded concomitantly at each position. We used the (110) chitin reflection in order to localize the cuticle and the Zn XRF signal for the localization of the cells. The intersect of the two signals marks the location of the assembly zone (Fig. 4A and C).

**Fig. 4.**
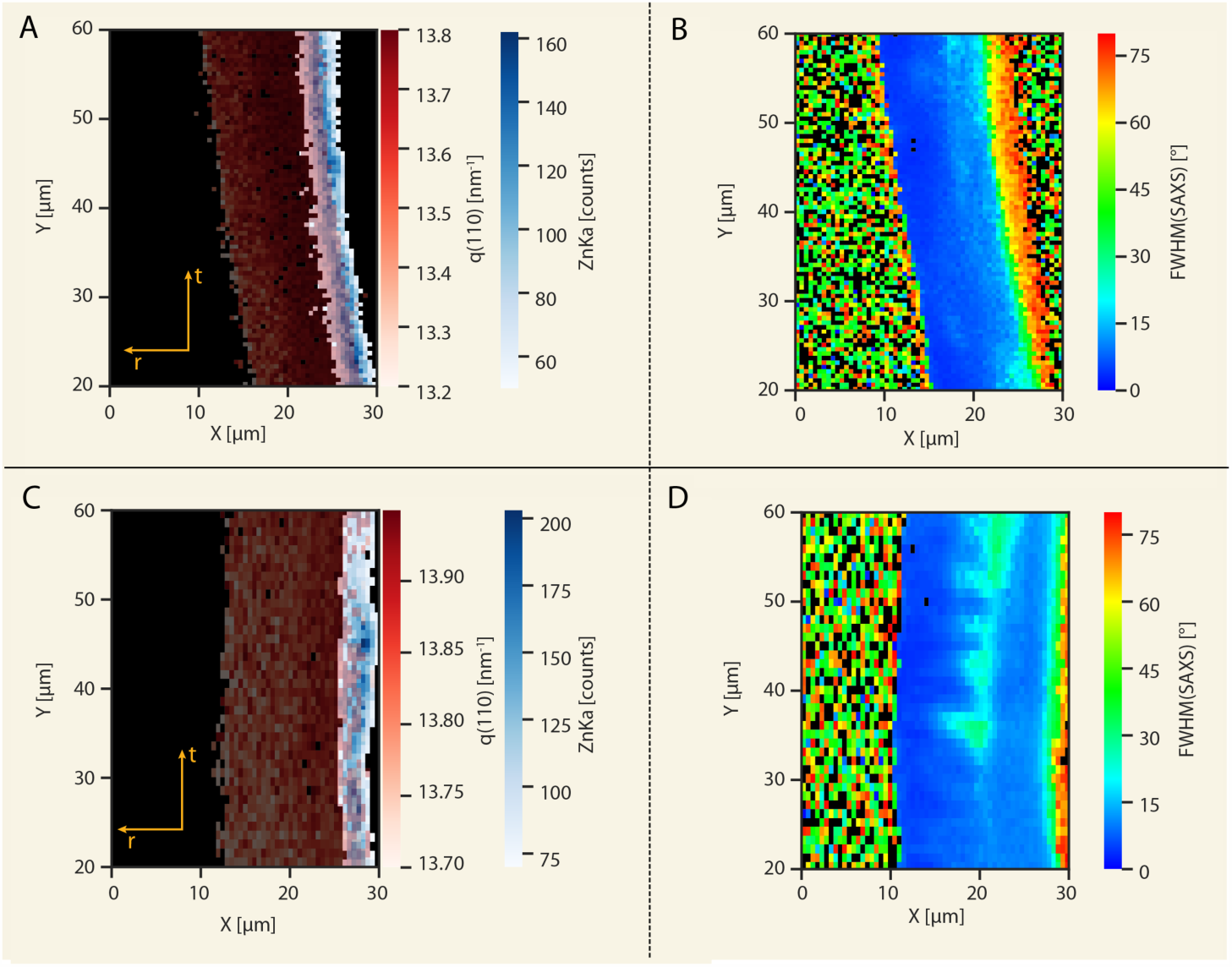
Scanning X-ray scattering and fluorescence results of locust tibia cross-sections. (A, C) Overlay of a map plotting the scattering vector (q) position of the (110) chitin reflection, and the Zn Kα X-ray fluorescence signal originating from the epidermal cells. The overlap between the two signals marks the assembly zone in (A) *Night* and (C) *Day* samples. (B, D) Heatmaps of the SAXS peak width, representing the degree of fiber alignment in (B) *Night* and (D) *Day* samples. The increased width of the SAXS peak in the assembly-zone indicates decreased fiber alignment and lower fiber compactization.

Small angle X-ray scattering (SAXS) provides information about the spatial nano-scale organization of the chitin-protein fibers within the assembly zone and within the cuticle, in the section plane. The SAXS intensity was integrated radially and plotted as a function of the azimuth angle (Supplementary Information Fig. 8). The width of the peaks in this plot is related to the degree of fiber alignment with respect to each other within the *(rt)* plane (as defined in Fig. 1A). The SAXS peak-width (FWHM, in units of degrees, Supplementary Information Fig. 8) is mapped in Fig. 4B and D across a tibia cross-section in *Night* and *Day* samples, respectively. In both cases the SAXS peak width at the assembly zone is broader than the peak width in regions within the cuticle bulk. This is supportive of a compactization process occurring after deposition as discussed above.

X-ray diffraction (XRD) provides information about the molecular structure, the respective organization, and the interaction of chitin with the cuticular proteins(23, 24). We have recently shown that co-ordering of chitin and proteins along the b-crystallographic direction of chitin, in the tarsal-tendon of the spider *Cupiennius salei*, gives rise to a shift in the (020) chitin reflection due to coherent scattering (24). In addition, protein-specific reflections with d-spacing equal to 0.47 nm, characteristic of intra-sheet inter-strand spacing in cross beta-sheet motifs were observed along the meridian in 2D fiber diffraction patters of spider tendons. In order to best identify protein-related reflections in the locust tibia, we reared animals in 24 h light settings. In these conditions, the endocuticle is made only of unidirectional layers (17). The fiber diffraction profiles obtained from longitudinal sections (∼70 µm thick) of such samples show a shift in the (020) reflection of chitin as well as several protein-related reflections along the meridian, at d = 0.47 nm (as observed in the spider tendon diffraction pattern) and at d = 0.43 nm (Supplementary Information Fig. 9). Inter-strand spacing of 0.47 nm is common for beta-sheet structures as in amyloid structures, whereas spacing of 0.43 nm is common for the inter-strand spacing in silks (25–27). Other unassigned reflections with d-spacing of 0.38 nm and 0.77 nm are observed along with additional reflections in the small angle region with d-spacing of 1.5 nm and 3.4 nm (the latter already noted by *Rudall et al*. for the cuticle of various insects species (28)). Based on these results we suggest that at least a large portion of the cuticular proteins contains cross-beta structures where the beta-strand orientation is roughly perpendicular to the chitin fiber long axis in unidirectional cuticle. Although individual protein reflections are not visible in helicoidal cuticles due to their low intensity and their overlap with chitin reflections, the presence of the ordered proteins is evident by apparent peak broadening of the chitin reflections as well as the shift in the (020) reflection.

Based on the structural information gained from unidirectional oriented cuticle, we used scanning X-ray diffraction to study the local chitin-protein molecular organization within different cuticular layers as well as within the assembly zone. Here, we used cross-sections of the tibia, such that the fiber orientation in unidirectional layers is parallel to the X-ray beam (similar to SAXS experiment above). In this orientation we identify an additional protein reflection with d-spacing of 1.1 ± 0.05 nm (q = 5.5 nm^-1^). Reflections at these positions are often assigned to inter-sheet spacing in stacking beta sheets when they are accompanied with orthogonal intra-sheet reflections like the ones observed here (d = 0.47 nm). This result is in good agreement with previous prediction of double layer of cross beta-sheet stacking(24). Note that according to the theory(24), when coherent chitin/protein scattering occurs, a separate protein reflection is unexpected. That this reflection is observed suggests that not all the proteins share a coherent interface with the chitin crystallites and/or that coherence occurs only along the chitin b-direction, while the proteins surround the chitin in all directions in the *ab* plane.

The (020) reflection of chitin and the protein inter-sheet reflection were peak-fitted for each diffraction profile within the scanned region. Mapping the intensity of the (020) peak (Fig. 5A), allows assigning regions of unidirectional organization (high peak intensity) produced during the day, vs. regions of helicoidal organization (low peak intensity) produced at night. The peak position of the (020) reflection is shifted in the cuticle to lower q values (q = 6.35 nm^-1^ to 6.4 nm^-1^), with respect to pure chitin (q = 6.61 nm^-1^) as seen before in samples from locusts grown in all-day conditions. In the assembly zone, however, this reflection is closer (q ∼ 6.5 nm^-1^) to pure chitin (Fig. 5B), suggesting that in this region the proteins and chitin are not yet co-ordered to produce a coherent diffraction interference. A map of the chitin to protein ratio based on the integrated intensity of the protein peak at 5.5 nm^-1^ and the (020) peak (Fig. 5C), as well as the azimuthal integration of 2D patterns taken from the assembly zone (Fig. 5D), show that the inter-sheet protein peak has lower intensity in the assembly zone relative to the cuticle (in both unidirectional and helicoidal regions).

**Fig. 5.**
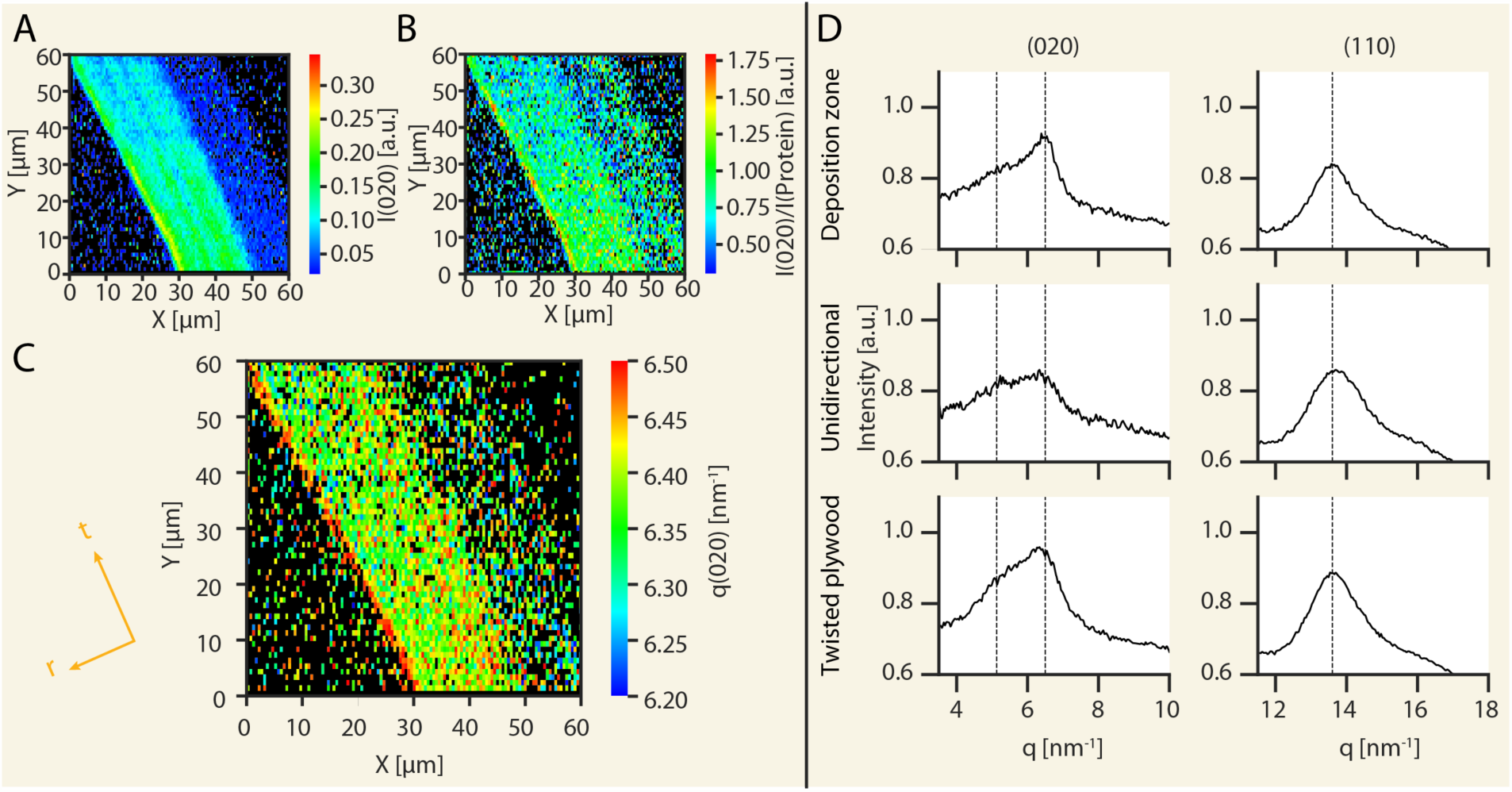
Scanning X-ray diffraction measurements of locust tibia cross-sections. (A) Intensity variation of the (020) reflection allows identification of "*Day*” (high I_(020)_) and "*Night*” (low I_(020)_) regions. (B) Peak position (q) of the (020) reflection. The q(020) position is shifted in the cuticle relative to the assembly zone. (C) Integrated intensity ratio between the chitin reflection (020) and the protein reflection (q∼5.5 nm^-1^). The ratio is increased in the assembly zone relative to the cuticle. (D) Averaged azimuthally integrated X-ray diffraction data showing the (020) and (110) reflections from the assembly zone (upper panel), the unidirectional *Day* regions (middle panel) and the helicoidal *Night* regions in the cuticle.

Increased protein diffraction in the cuticle accompanied with the (020) peak shift (Fig. 5) suggests chitin-protein co-assembly. Thus, compactization, as observed from SAXS analysis, and co-alignment of chitin and protein, as judged from the XRD analysis, are co-localized implying that co-assembly of the protein and chitin may be the driving force for fiber re-orientation. We note, that the phase diagram of chitin in water is extensively explored. Unfortunately, we cannot relate our findings directly to the lyotropic behavior of chitin as determined from *in-vitro* experiments in aqueous solutions. Indeed it is well known that chitin forms cholesteric LC phases with different helicoidal pitch depending on e.g. pH, ionic strength and crystallite size(29–31). However, the physico-chemical conditions in the assembly zone are to date unknown and the effect of chitin-binding proteins on chitin assembly has not been systematically addressed experimentally *in vitro*.

**In summary**, our results suggest that dynamic apical cell surface structures in the cuticular epidermal layer determine the boundary conditions for fiber self-assembly by secreting the fibers into the assembly zone in pre-determined orientations. During the night, when helicoidal fiber arrangement is formed, the fiber orientations are discrete, resembling the microvilli spatial organization, whereas, when parallel fiber arrangement is produced during the day, the fiber orientation in the assembly zone, is parallel to the elongated microvilli ridges structures. The final orientation of the fibers is achieved by concomitant chitin-protein co-ordering and compactization of the cuticle resulting in a stable helicoidal or unidirectional structure in the endocuticle. We note that the helicoidal organization could, theoretically, be produced by employing similar cell surface structure as seen in light conditions and rotating the orientation of the ridges between deposition cycles to produce the rotated plywood structure. This is however not the case, since the apical cell surface structures differ significantly in microvilli morphology between light and dark conditions. This leads us to suggest varying levels of cellular control on the assembly process. Here, the unidirectional arrangement is under direct cellular control on fiber orientation whereas the helicoidal structure is obtained away from the apical cell surface. Based on our X-ray data we propose that chitin-protein co-assembly and protein-protein interactions are the driving forces for the process. Our results provide direct evidence for the two-model system and the self-assembly hypotheses put forward by Neville(16), Giraud-Guille and Bouligand(15, 32) based on the architecture of the cuticle in its final form and its resemblance to liquid-crystal geometries observed *in vitro*. The observations and the methodology presented here could be relevant for a variety of biological extra-cellular matrices where self-assembly and cellular controlled mechanistic questions are yet unresolved.

## Methods

First, fourth and fifth instar specimen of *Locusta migratoria* were obtained from Reptilienkosmos (www.reptilienkosmos.de). The animals were reared in 12/12 light cycle condition or in 24 h light and 36 °C/26 °C or 36 °C conditions, respectively. The light source was a Mini light strip LED (Lucky Reptile) emitting a daylight spectrum at 200 lm. Tibiae samples were obtained 2 days to 2 weeks after ecdysis into the next stage (next instar or adult). *Day* samples were obtained during the light phase of the illumination cycle whereas *Night* samples were obtained during its dark phase.

### FIB/SEM sample preparation

Chemical fixation: Three biological replicates of hind tibia of both fifth instar and adult *L. migratoria* were collected during light and dark phase of the illumination cycle, 3 days after ecdysis. The entire hind tibia was submerged into 2.5% (w/v) glutaraldehyde and 2% (w/v) paraformaldehyde in 0.1 M cacodylate buffer (pH 7.4) and incubated for 4 h at room temperature. Samples were then washed 5 times for 10 min in 0.1 M cacodylate buffer and incubated for 2 h in 2% (w/v) osmium tetroxide. The samples were washed again in 0.1 M cacodylate buffer 5 times for 10 min each and dehydrated in a series of ascending concentrations of acetone (30%, 50%, 70%, 90%, 100% 2x) with 10 min for each step. Tissues were then infiltrated with Durcupan/Epon epoxy resin (Matsko and Mueller 2004) and cured in an oven at 60 °C for 48 h. Cross-sections of the upper region of distal part of hind tibia were prepared (see Supplementary Information Fig. 10), coated with 5 nm carbon and 5 nm platinum and imaged using Focused Ion Beam / Scanning Electron Microscope (FIB/SEM).

Cryo-fixation: At least 3 biological replicates of hind tibia of third, fourth and fifth instar and adult *L. migratoria* were collected during light and dark photoperiod. Thin (fifth instar and adult stage animals) or thick (third and fourth instar stage animals) cross-sections of the upper region of distal part of hind tibia were prepared (see Supplementary Information Fig. 10) with tibia submerged into standard locust saline solution (36). We did not observe structural differences in apical cell surface or in the assembly zone that could be related to the developmental stage of sampled animal. Cross-sections were vitrified using high-pressure freezing machine (HM100, Leica) in 20 mM dextran solution for cryo-protection. Vitrified samples were freeze-substituted with a fixation and staining cocktail (1% osmium tetroxide, 0.1% uranyl acetate, 0.5% glutaraldehyde, 1.5% water) in acetone for 2 days at −85 °C using automatic-freeze substitution machine (AFS2, Leica). The samples were warmed up to room temperature over 1 day and embedded as described above. Polymerized resin blocks were further polished to expose the cross-sections for block-surface imaging. After coating the surface with carbon and platinum as described above, samples were imaged using FIB/SEM. In chemically fixed samples, the formed fibers in the assembly zone cannot be resolved due to excessive cross-linking leading to over-staining. We therefore used the geometry of the pore canals (Supplementary Information Fig. 2 and 3) to determine the fiber orientation in the last deposited layers: the pore-canals are channels running throughout the cuticle containing cellular processes, characterized by their almond shape cross-section. When the pore canals are passing through lamellate helicoidal cuticle the cross section shows a continuous rotation, which is absent in non-lamellate unidirectional cuticle architecture(16).

### FIB/SEM image serial imaging (Crossbeam 540, Zeiss)

For chemically fixed dataset representing *Night* samples 538 serial electron micrographs were acquired in *rt* plane with voxel size (13.55 × 13.55 × 17.5) nm^3^. For dataset representing chemically fixed *Day* samples 393 serial electron micrographs were acquired in *rt* plane with voxel size 12.82 × 12.82 × 17.5 nm. In both cases, SEM imaging was performed at 2 kV acceleration voltage and 1 nA probe current using secondary electron detector while slices were generated using 300 pA FIB probe at 30 kV accelerating voltage.

For dataset representing cryo-fixed samples the number of slices ranged from around 800 to 2270 and voxel size from (5.28 × 5.28 × 10.5) m^3^ to (8.1 × 8.1 × 21) nm^3^. In all cases, SEM imaging was performed at 2 kV acceleration voltage and 1 nA probe current using secondary electron detector while slices were generated using 300 pA FIB probe at 30 kV accelerating voltage.

### Transmission Electron Detector (TED) imaging

TED imaging was performed inside the FIB/SEM chamber using STEM4A detector. 100 nm thin sections of cryo-fixed samples were prepared using a Leica UC-6 microtome. The sections were mounted on copper grids and imaged with 20 kV acceleration voltage and 300 pA probe current using high resolution column mode.

### FIB/SEM Image analysis

Image processing and segmentation of data cubes were performed as follows: First, images belonging to each stack were registered taking advantage of the Fourier shift theorem. We calculated the displacement vector of each image to the previous one (taking the first image of the stack as a reference), ***d***_*i*_, using the phaseCorrelate function of OpenCV (v 2.4.7). Then, the shift of each image with respect to the first one, ***D***_*i*_, was calculated as ***D***_*i*_ = S_i_***d***_*i*_. In many cases, curtaining artifacts were visible in the stacks, giving rise to vertical contrast modulation (stripes). To correct for this, we followed the approach described in a recent paper (37), where the depth of the wavelet transform, the wavelet family and the width of the Gaussian filter were visually optimized to obtain a corrected stack with the least contrast and information loss and the largest stripe removal. Finally, to improve the signal to noise ratio, the image stacks were denoised using the skimage.restoration local denoising filters of the scikit-image python library (v. 0.14.0). Again, the filtering algorithms and parameters were chosen by visually optimizing the corrected images to obtain the largest noise removal with the lowest information loss.

Segmentation of structures of interest was performed semi automatically using a combination of Amira 3D (FEI, USA) and a custom python code implementing state of the art 2D and 3D machine learning methods using Keras and GPU accelerated Tensorflow. For the 2D segmentation between 20 and 30 representative 256 × 256 regions of the image stack were sparsely annotated using Amira 3D. The images and ground truth labels were augmented using the Keras ImageDataGenerator class by allowing a rotation and shearing. The augmented images were used to train a U-net like network which consists, for the contracting path, of the repeated application of 3×3 convolutions (padding = same) followed by a ReLU activation and 2 × 2 max pooling for downsampling. For the expanding path, every step consists of a 2 × 2 upconvolution concatenated with the corresponding layer of the contracting path and the repeated application of 3×3 convolutions followed by a ReLU activation. A dropout of 0.25 after each max pooling and upconvolution operations was applied. A total of 4 max pooling and upconvolutions were used (depth of the network = 4). On the final layer, a sigmoidal activation was employed.

The input images and the corresponding ground truth labels were used to train the network on a workstation with a single Nvidia Quadro M5000 GPU with 8GB of memory with the adam optimization algoritm, a learning rate of 1e-4 and binary cross entropy as a loss function. Once the loss dropped below 1%, the training was stopped and the network was used to infer the labels on the whole image stack. Although the network would predict the micovilli with a high accuracy, in many of the images would contain a small amount of predicted labels not coinciding with the objects of interests. These were manually deleted from the image stack using Amira 3D.

For the 3D segmentation, a similar workflow was used: first, 2 representative sub volumes (512 × 512 × 512) px^3^ of the image stack were selected and densely manually annotated using Amira 3D. From these 2 volumes, we extracted randomly 30 smaller volumes (92 × 92 × 92) px^3^. Each of these 30 volumes was randomly transformed 3 times to generate 3 ‘augmented’ volumes using one of the following transformation: identity, reflection with respect to the x or y or z axis, 10% zoom in or out and 10% dilation or compression on a random axis (x, y or z). The generated augmented volumes were randomly cut to a final size of (64 × 64 × 64) px^3^ not containing boundary artifacts associated with the transformation (like for zooming out and compressing). The final 90 volumes (64 × 64 × 64) px^3^ were used to train a 3D network with an architecture slightly modified with respect to the one used for the 2D segmentation. The basics building blocks of the network remained unchanged, but we used convolution kernels of (5 × 5 × 5) px^3^, a network’s depth of 2 instead of 4 and the layers in the contracting and expanding path were not concatenated.

The network was trained with the same hardware and software implementation as in two dimensions. In this case, the training was stopped once the loss reached a value of 0.05 and the network used to infer the whole volume. Because of the limitation of the GPU memory size (8Gb), the inference was run by tiling the whole volume in tiles of (128 × 128 × 128) px^3^ overlapping for half of their size. For each tile, only the inferred labels in the central (64 × 64 × 64) px^3^ were used to generate the final predictions on the original image stack. Because the training sub-volumes were taken from the image stack the inference was run on, the quality of the predictions in those regions was superior to the others. Nevertheless, large portions of the stack neighboring those selected for the training exhibited also accurate predictions. Also in this case, the predicted labels were checked with Amira 3D and the wrongly predicted labels were manually deleted. The Drishti volume exploration and representation tool (v.2.6.5) was used for volume rendering (38). Orientation maps and distributions were generated with Orientation J plugin of Fiji (http://bigwww.epfl.ch/demo/orientation/). Three-dimensional Fourier transforms of the stack of the labelled microvilli were calculated using scipy (v 1.3.0). Azimuthal averages of the 2D Fourier transforms were calculated using a custom script in python.

### SAXS/WAXS Sample preparation

The distal tibia part of *L. migratoria* adult animals reared in 12 h dark/ 12 h light cycle or 24 h light were severed from the animals several days after ecdysis, embedded in O.C.T (VWR Chemicals, Radnor, PA, USA) and rapidly frozen using liquid nitrogen and a silicone mold. The frozen blocks were sectioned using a HM 560 CryoStar Cryostat (Thermo Fisher Scientific, Waltham, MA, USA) with Surgipath DB80 LX blades (Leica, Wetzlar, Germany) (Sample −15 °C, blade −11 °C). For the 12 h dark/ 12 h light samples cross sections of 20 µm and 30 µm thickness were prepared for SAXS and WAXS measurements, respectively. In case of the 24 h light samples, longitudinal sections of 70 µm in thickness were cut. The sections were thawed, rinsed with water and transferred to a SiN membrane (Silson, Southam, UK) (3 × 3 window array (each (5 × 5) mm^2^, 1 µm thick), frame thickness 200 µm (23.5 × 23.5) mm^2^ total size) for XRD/XRF measurement.

### SAXS/XRD and XRF mapping

X-ray fluorescence and scattering experiments on *L. migratoria* tibia sections were performed at the microXAS – X05LA beamline at the Swiss Light Source (SLS) synchrotron radiation facility (Paul Scherrer Institute, Villigen, Switzerland). The X-ray beam was defined to 15.2 keV (0.816 Å) using a Si(111) monochromator and focused to (1 × 1) µm^2^ using KB mirrors. XRD data was obtained using a 2D Dectris Eiger 9M detector ((2070 × 2167) px^2^) behind the sample, while a single element Si(Li) detector (Ketek, Munich, Germany) was used for XRF measurements located perpendicular to the beam main axis. Samples were mounted at a sample-to-detector distance of around 250 mm on a rotating x, y, z-stage. XRD and XRF signals were recorded simultaneously with an integration time of 1 s. The scanning step size in y direction was 1 µm, in x direction 0.5 µm. Quartz powder was used to calibrate the sample to detector distance.

XRD data from 24 h day sample was obtained at the µSpot beamline, station of the l-Spot beam line at the synchrotron BESSY II (Helmholtz Center, Berlin, Germany). The X-ray energy of 15 keV was defined by a multilayered monochromator. The incident X-ray beam was focused on the sample by a toroidal mirror, and the final beam size was defined by a pinhole of 10 µm diameter placed in front of the sample. WAXS data were collected using a large-area 2D detector (MarMosaic 225, Mar USA Evanston, USA), situated approximately 300 mm behind the sample.

Calibration, integration and data analysis of the 2D diffraction patterns were performed using the software DPDAK (39). DPDAK was also used for background removal and peak-fitting the XRD patterns. For each peak, several fits were performed where the fitting parameters (q position, intensity and peak width) were first varied and then fixed. We used Lorenzian peak-shape for fitting in q space (neglecting micro-strain contribution) and Gaussian peak-shape for fitting in azimuthal space. OriginPro 2015 software was used for averaging of 1 d diffraction patterns. 2D heatmaps were plotted using the matplotlib library in Spyder 3.3.2 from the Anaconda package (40).

### Confocal laser scanning microscopy (CLSM)

Sections were prepared as indicated above using a cryo microtome, and stained with Nile red and DAPI in standard locust saline solution (36) for 1 h or with Direct Yellow 96 (Sigma Aldrich) in Standard Locust Saline solution for 1 h. Images were acquired using a SP-8 laser scanning confocal microscope (Leica) equipped with a 63x water immersion objective (NA = 1.2). For excitation of DAPI and Direct Yellow 96, a multiphoton laser with a wavelength of 780 nm was used and the signal was recorded by a HyD detector with a bandpass filter set to 400 nm to 480 nm and 540 to 610 nm, respectively. For Nile Red a DPS S 561 laser emitting light of 561 nm was used for excitation together with a HyD detector with a bandpass filter of 570 nm to 620 nm for detection of the fluorescence signal.

## Acknowledgements

We appreciate the help of Birgit Schonert from the Max Planck Institute of Colloids and Interfaces with sample preparation. We thank Prof. Thomas Stach (Humboldt University, Berlin) Dr. Katja Höflich and Dr. Holger Kropf (Helmholtz-Zentrum Berlin für Materialien und Energie, Berlin, Germany) and Dr. Igor Zlotnikov (B CUBE, Technical University Dresden, Dresden) for assistance with TEM and FIB/SEM imaging. Thank you to Sebastian Wieser, Lisa Schoop and Romy Angermann for help with animal maintenance and sampling. We are grateful to Dr. Changhao Li and Dr. Stefan Siegel for beamtime assistance during the measurements at BESSY II, Helmholtz-Zentrum Berlin (HZB), Germany and to Dr. Grolimund Daniel and Dr. Dario Ferreira Sanchez from the Swiss LightSource, Paul Scherrer Institute (PSI), Vilingen, Swizerland for invaluable assistance during beamtime at the microXAS - X05LA beamline. We acknowledge HZB, Germany and PSI, Villigen, Switzerland for the allocation of synchrotron radiation beam time at beamlines µSpot at Bessy II and microXAS-X05LA of the SLS, respectively. A special thank you is given to Prof. Friedrich Barth, University of Vienna, Vienna, Austria, Prof. Nadine Nassif, Sorbonne Universités, CNRS, Paris, France and Dr. Yoshiharu Nishiyama, University Grenoble Alpes, CNRS, CERMAV, Grenoble, France for fruitful and inspiring discussions.

## Data and materials availability

All data, code and materials used is available upon request

