## Supplementary data for "Epidermal cell surface structure and chitin-protein co-assembly determine fiber architecture in the Locust cuticle"

**Supplementary Information**

**Supplementary Information Fig. 1. FIB-SEM slices of cryo-fixed locust tibia showing microvilli structures**.

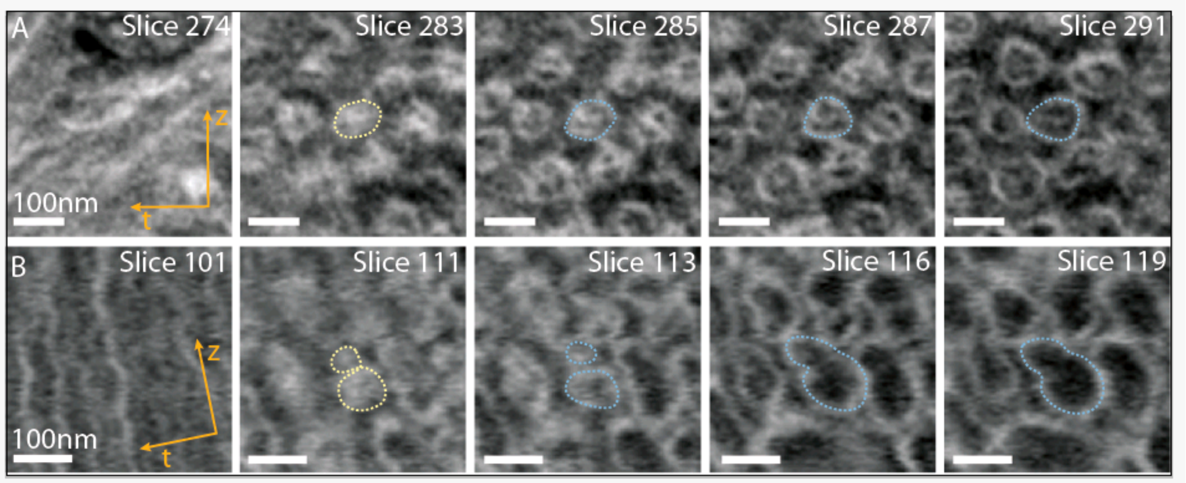


(A) *Night* (B) *Day*. The plaques are indicated with a light yellow dotted line and the microvilli base with a light blue dotted line. The slices are presented clockwise from the cuticle to the cells, i.e. along *(r)* direction.

**Supplementary Information Fig. 2. FIB/SEM slices of chemically fixed locust tibia**.

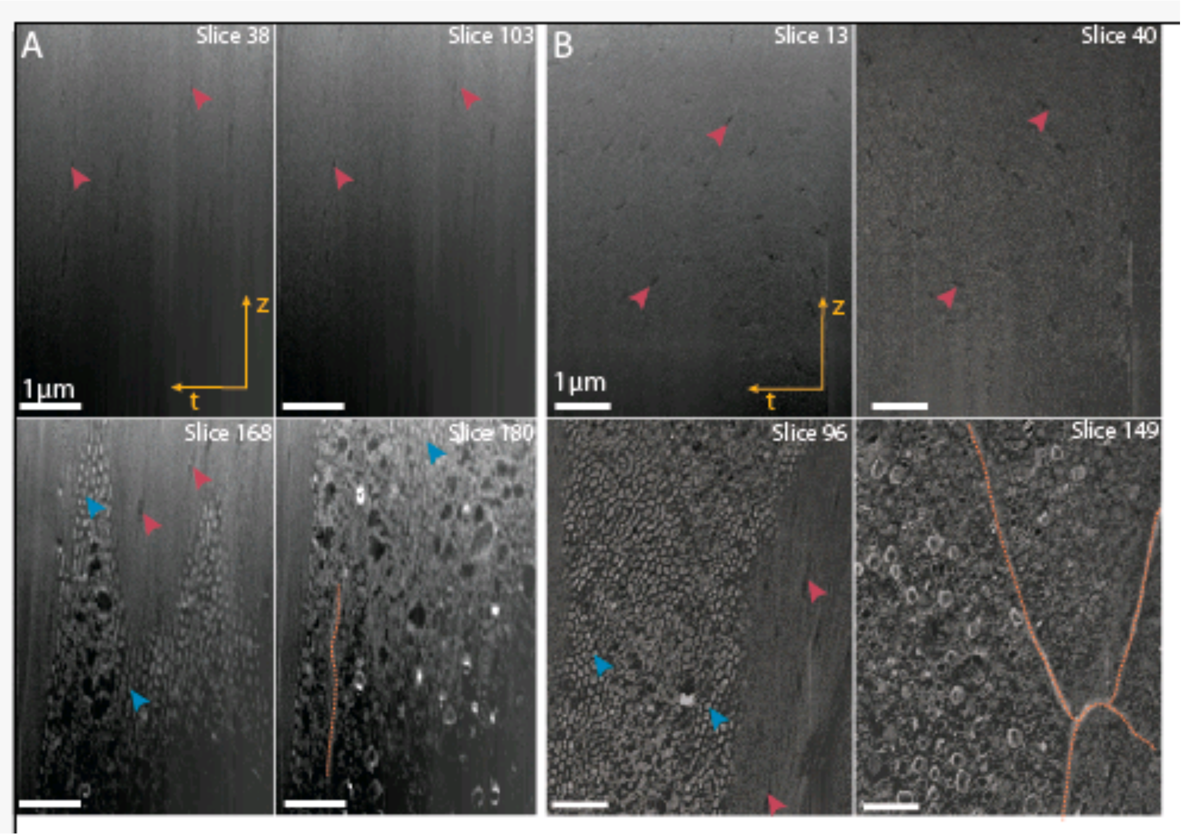


(A) *Night* sample. (B) *Day* sample. Marked: Pore canals in the cuticle (magenta arrowheads), cell borders (dotted orange lines), apical cell protrusions (blue arrowheads). The slices are presented clockwise from the cuticle to the cells, i.e. along *(r)* direction.

**Supplementary Information Fig. 3. Apical cell-surface structures in *Night* and *Day* samples. Reconstruction and quantification of 3D FIB/SEM data obtained from chemically fixed locust tibiae samples.**

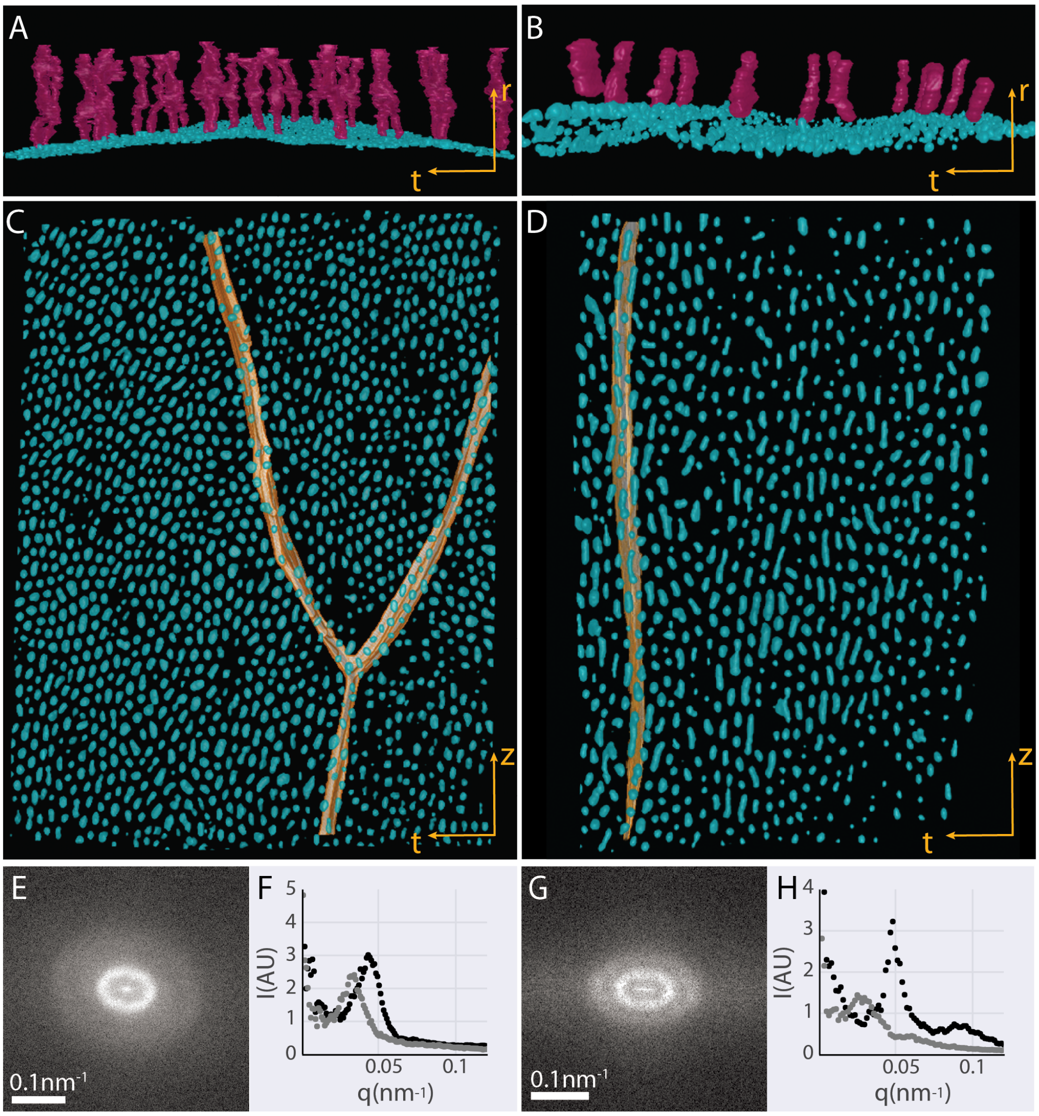


(A, B) Volume rendering of the segmented apical protrusions at the surface of the epidermal cells (cyan) and the pore canals (magenta) in *Night* (A) and *Day* (B) samples. (C, D) 'Top-view' of the rendered volume of the segmented apical surface of the epidermal cells (cyan) and the lateral cell membrane (orange) in *Night* (C) and *Day* (D) samples. Note that the respective apical protrusions organization is continuous across multiple cells. (E, G) 3D Fourier-Transformation (FT) of the segmented volumes representing the epidermal cell surfaces in *Night* (E) and *Day* (G) samples. (F, H)

Azimuthal partial integration of the FT pattern in (E) and (F), respectively in the meridional (light grey) and equatorial (dark grey) directions, showing increased isotropy and reduced long-range order correlation of the apical protrusions in *Night* vs. *Day* samples.

**Supplementary Information Fig. 4. TED micrographs of the assembly-zone**.

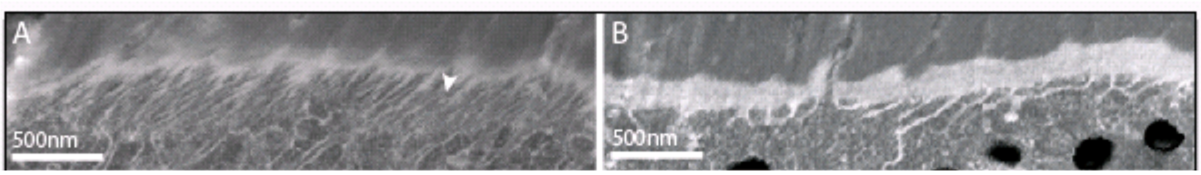


Comparison of the assembly zone in areas where microvilli are present (A) or absent (B). STED micrographs obtained from a 100 nm thick section, sense structures are bright, as in FIB/SEM micrographs. Arrowhead in (A) points to the smallest/thinnest resolved fiber.

**Supplementary Information Fig. 5. FIB/SEM slice series from regions in a *Day* sample lacking microvilli**.

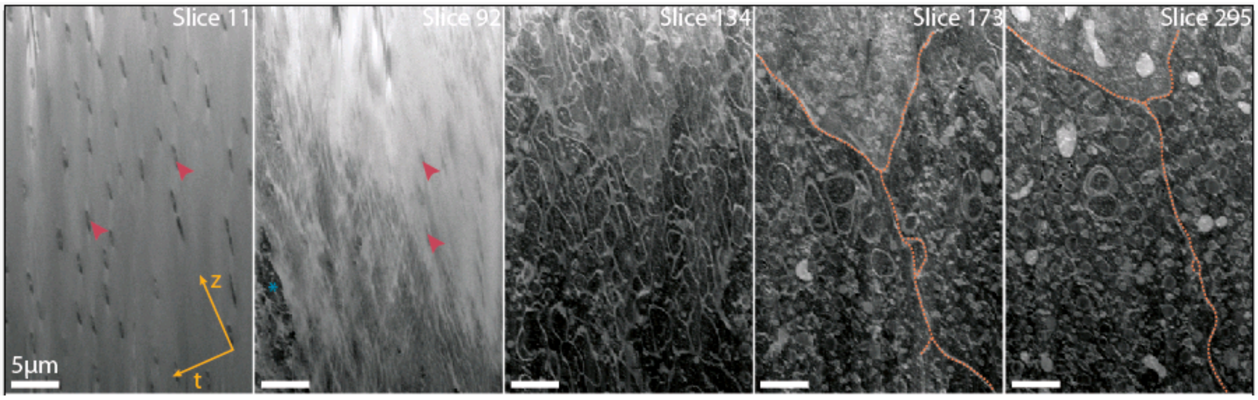


Magenta arrows point to pore-canals. Dashed orange lines indicate lateral cell membranes. Reconstructed volumes are presented in Fig 3 in the main-text.

**Supplementary Information Fig. 6. Segmentation of FIB/SEM data**

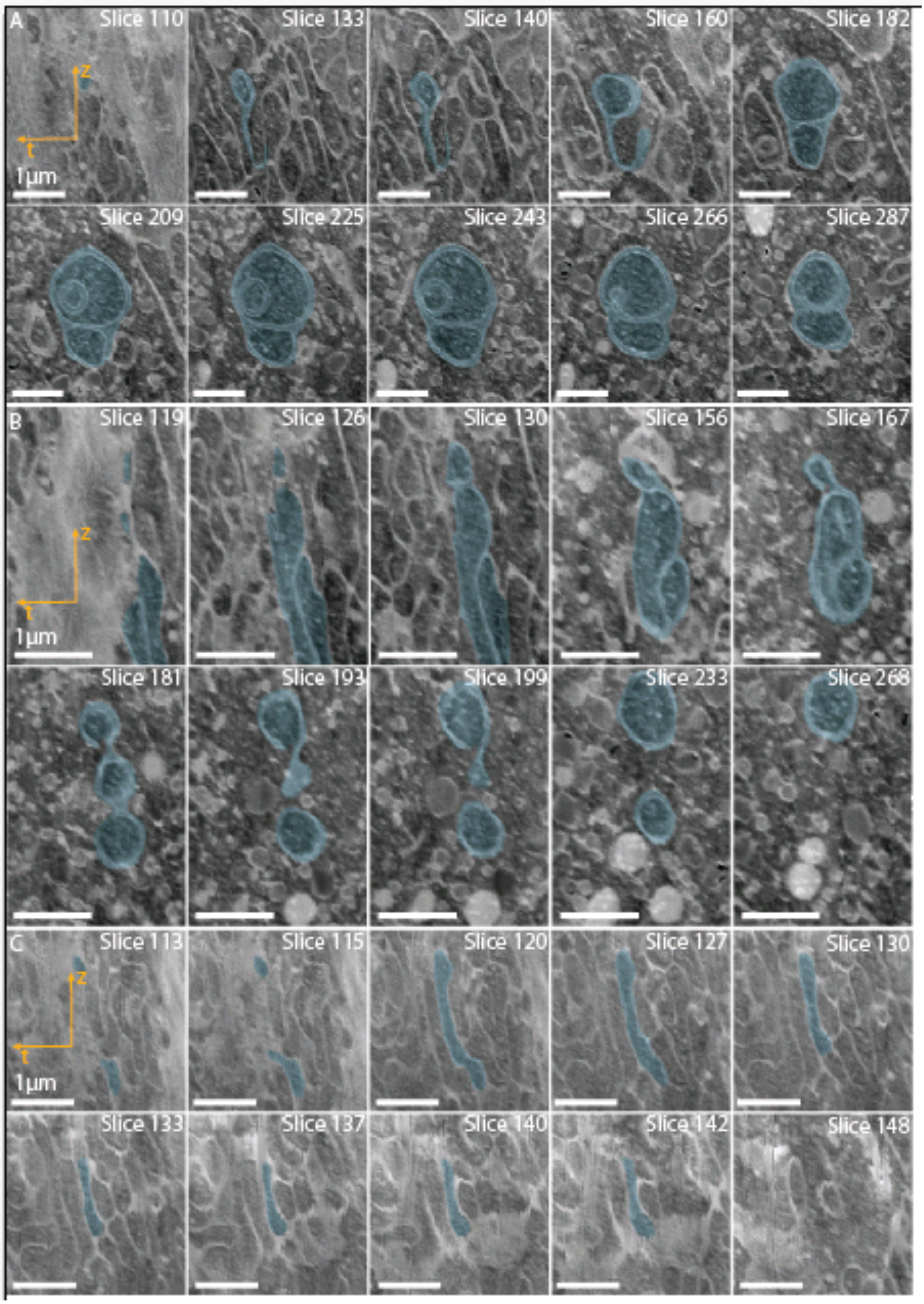

FIB/SEM slice series from regions in a *Day* sample presented in 3D volume rendering in Fig. 3. E, F and g correspond to features depicted in Fig. 3 in main text.

**Supplementary Information Fig. 7 FIB/SEM data of epidermal cell apical surfaces.**

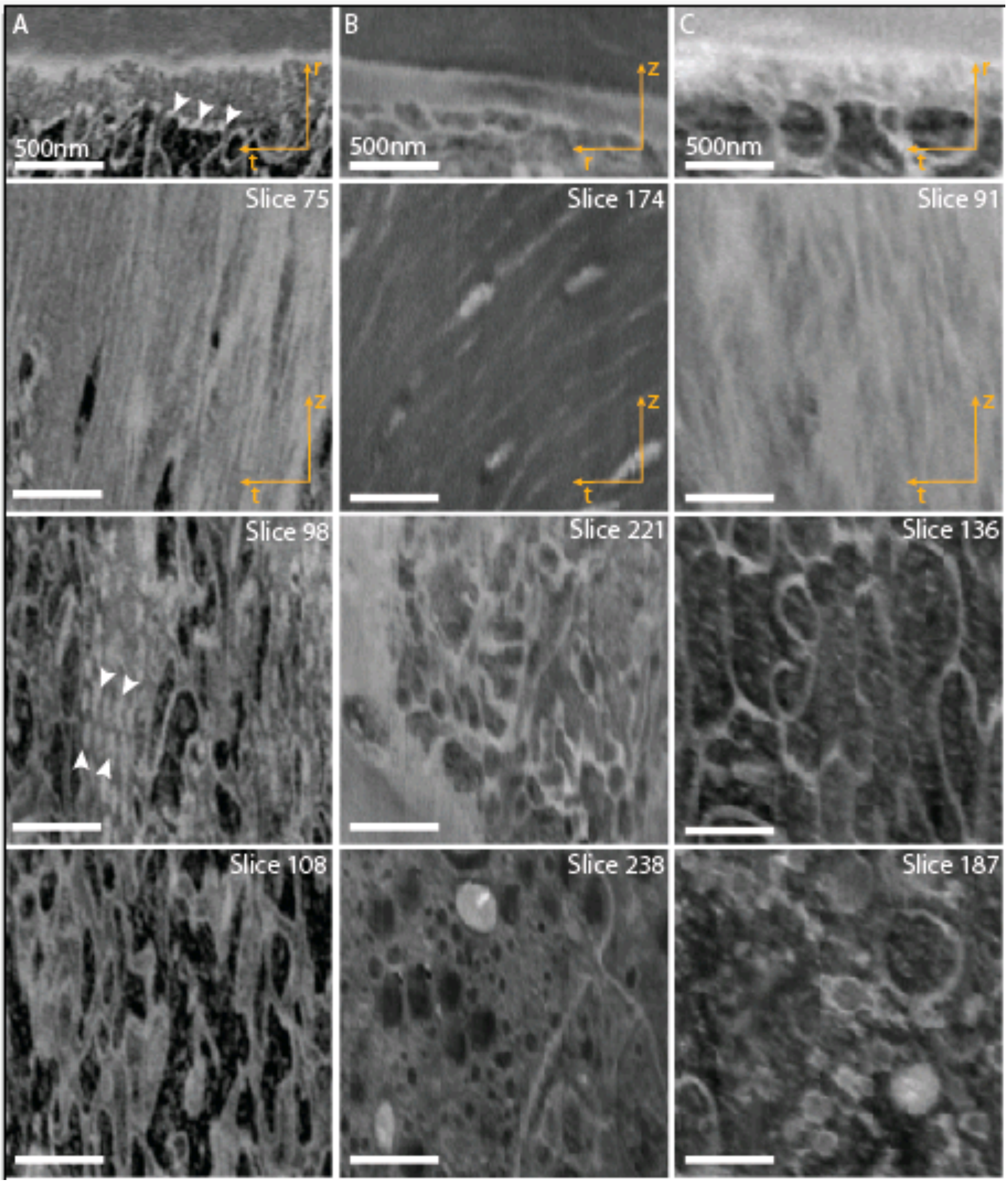


Data acquired in regions in *Day* (A, C) and *Night* (B) samples where microvilli are partially (A) or fully absent (B, C). Upper panel shows the *(rt)* or *(rz)* planes, followed by slice series along the r direction displaying the *(tz)* planes. Plaque-like structures in (A) are marked by white arrowheads.

**Supplementary Information Fig. 8. SAXS peak azimuthal width.**

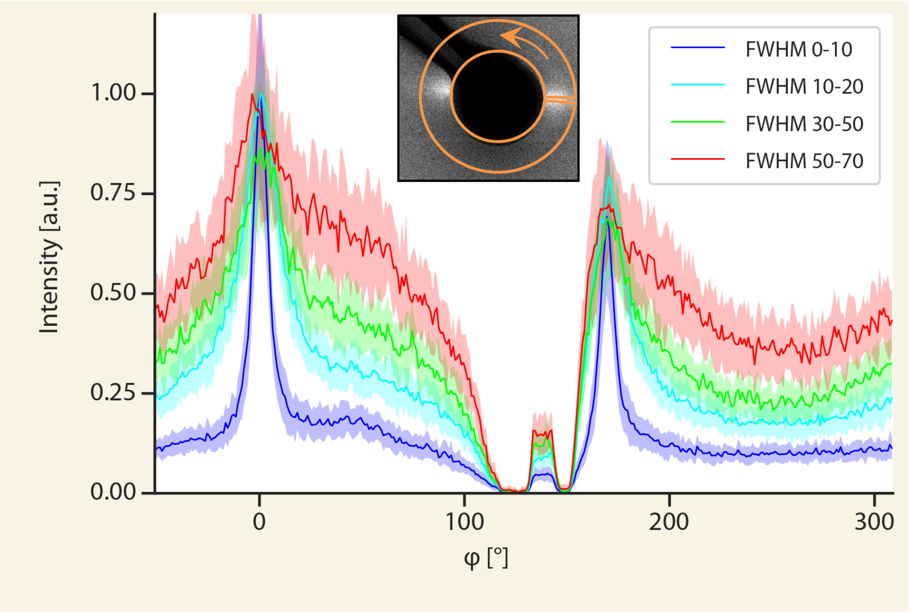


Radial integration of the SAXS patterns obtained from various regions in the cuticle and assembly zone on a locust tibia section. The widths of the peaks (FWHM) are displayed in Fig. 4b, d. Inset shows an example of a 2D SAXS pattern from the cuticle region. The orange arrow marks the integration direction, and the orange contour shows the radial and azimuthal range used for the integration. Single profiles are averaged from multiple spectra in the given range with the standard deviation displayed in light color around it.

**Supplementary Information Fig. 9. WAXS data of locust tibia grown in 24 hr light cycle.**

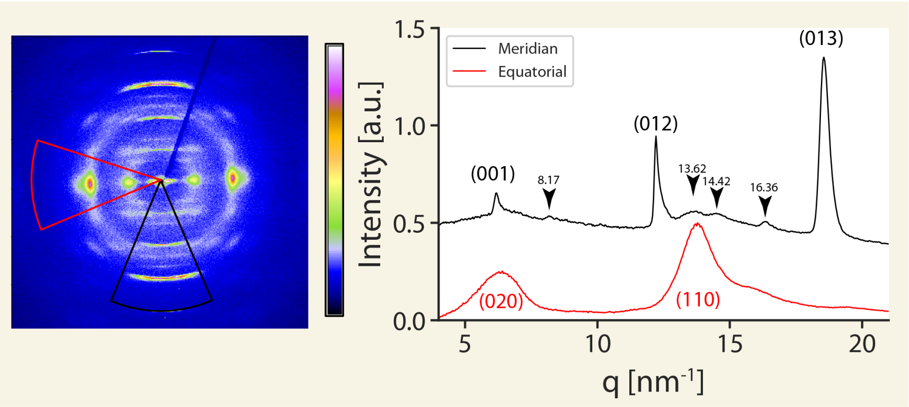

2D XRD pattern and azimuthally integrated 1D XRD profiles in the equatorial direction (red line, red cake on 2D pattern) and meridian (black line, black cake on 2D pattern), obtained from the endocuticle of a locust reared in 24 hours day conditions. Black arrows point to protein diffraction peaks at q = 8.17 $nm^{-1}$, 13.62 $nm^{-1}$, 14.42 $nm^{-1}$ and 16.36 $nm^{-1}$ corresponding to d spacing of 0.77 nm, 0.46 nm, 0.44 nm and 0.38 nm.

**Supplementary Information Fig. 10. Hind tibia of *L. migratoria* contains proximal and distal part**.

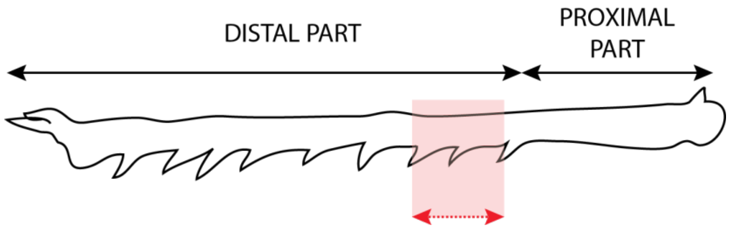


Area marked in red, upper region of distal part was sampled and analyzed in this study.
